# Activity-dependent changes in synaptic protein complex composition are consistent in different detergents despite differential solubility

**DOI:** 10.1101/576074

**Authors:** Jonathan D. Lautz, Edward P. Gniffke, Emily A. Brown, Karen B. Immendorf, Ryan D. Mendel, Stephen E.P. Smith

**Author notes:** Correspondence to: S.E.P.S.

## Abstract

At the post-synaptic density (PSD), large protein complexes dynamically form and dissociate in response to synaptic activity, comprising the biophysical basis for learning and memory. The use of detergents to both isolate the PSD fraction and release its membrane-associated proteins complicates studies of these activity-dependent protein interaction networks, because detergents can simultaneously disrupt the very interactions under study. Despite widespread recognition that different detergents yield different experimental results, the effect of detergent on activity-dependent synaptic protein complexes has not been rigorously examined. Here, we characterize the effect of three detergents commonly used to study synaptic proteins on activity-dependent protein interactions. We first demonstrate that SynGAP-containing interactions are more abundant in 1% Deoxycholate (DOC), while Shank-, Homer-and mGluR5-containing interactions are more abundant in 1% NP-40 or Triton. All interactions were detected preferentially in high molecular weight (HMW) complexes generated by size exclusion chromatography, although the detergent-specific abundance of proteins in HMW fractions did not correlate with the abundance of detected interactions. Activity-dependent changes in protein complexes were consistent across detergent types, suggesting that detergents do not isolate distinct protein pools with unique behaviors. However, detection of activity-dependent changes is more or less feasible in different detergents due to baseline solubility. Collectively, our results demonstrate that detergents affect the solubility of individual proteins, but activity-dependent changes in protein interactions, when detectable, are consistent across detergent types.

## Introduction

The postsynaptic density (PSD) is an electron dense specialization composed of multiprotein complexes, whose functions include mediating the apposition of pre-and post synaptic membranes, clustering glutamate receptors, and coupling the activation of receptors to downstream signaling cascades (1). Proteomic profiling of the PSD has revealed a highly complex and massive structure (1 GDa (2)), with the number of putative PSD proteins ranging from a few hundred to nearly two thousand (3-5). Scaffold proteins including the DLGs (e.g. PSD-95), SRC homology 3 and multiple Ankyrin repeats (Shank), Homer, and Synapse-associated protein associated protein (SAPAP) families are highly abundant (6, 7). Super resolution and electron microscopy have shown that these proteins are arranged in a layered organization, providing the basic structural framework for the PSD and serving as a molecular platform onto which other proteins are recruited (8-10). The most abundant scaffold protein, PSD-95, can bind to both NMDAR’s and AMPAR’s, either directly or indirectly, stabilizing the presence of these glutamate receptors at the PSD (7, 11). Through its PDZ domain, PSD-95 interacts with SAPAP, which in turn binds to the Shank family of proteins (12, 13). Shank and its binding partner Homer then form a polymerized, mesh-like second layer of the PSD, characterized by Shank dimers binding the EVH1 domain of Homer tetramers (13). Homer EVH1 domains can also bind type 1 mGluR receptors, IP3 receptors, TRPC channels, dynamin, and drebrin. Signaling enzymes such as Calcium/Calmodulin dependent Kinase II (CamKII) and the brain specific Ras GTPase, SynGAP are also highly abundant (14, 15), further increasing the complexity of the PSD (16, 17). Critically, the organization of the PSD is also highly dynamic. We and others have demonstrated that synaptic stimulation elicits dissociation of mGluR5-Homer-Shank scaffolds (18, 19) and dispersion of SynGAP from the PSD (18, 20-22), allowing for the recruitment and stabilization of AMPA receptors to the PSD and changes in synaptic strength (22, 23). It is this change in synaptic transmission efficacy that enables the PSD to regulate the flow of information between neurons and ultimately control the complex microcircuits to drive adaptive behaviors. Thus, dynamic synaptic protein co-associations at the PSD play a central role in synaptic plasticity, learning and memory.

The study of protein co-associations requires the optimization of detergent conditions such that proteins are solubilized, but co-associations are not disrupted. However, traditional biochemical analysis of the PSD has involved differential solubility in a series of detergents: Synaptosomes, or “P2” fractions, are solubilized in the non-ionic detergent Triton X-100 (Triton), and following high-speed centrifugation, the Triton-insoluble pellet is solubilized in the ionic, bile-acid deoxycholate (DOC) at a high pH (9.0) to yield a “PSD fraction” (24, 25). DOC belongs to a class of detergents often used for membrane disruption and extraction of membrane proteins, but being a strong detergent, DOC might also be expected to disrupt protein co-associations. (Figure 1). While many large protein complexes do survive DOC solubilization and have been extensively characterized by mass spectrometry and other approaches (7, 11, 26, 27), interactions that might be present at the native PSD but that do not survive DOC solubilization may remain biochemically uncharacterized. Recently, we and others have reported activity-dependent changes in protein co-associations involving presumably synaptic proteins using the relatively gentle, non-ionic detergent NP-40 (18, 19, 28, 29). For example, Ronesi et. al (26) showed that Homer and mGluR5 acutely dissociate in response to synaptic activity in a CAMKII-dependent manner. Using multiplex co-immunoprecipitation of lysates solubilized in NP-40, we reported the dissociation of 34 synaptic protein co-associations following acute stimulation of cultured neurons with NMDA, DHPG, or glutamate. However, because the PSD, as it is traditionally defined, is not soluble in NP40, it is unclear if these dissociations represent proteins localized to the PSD, or a group of peri-synaptic proteins. The Huber group showed that levels of mGluR5 found in a PSD fraction are also reduced following activity, suggesting direct relevance of the mGluR-Homer dissociation (29). Similarly, Frank et al. (11) reported that PSD-associated protein super-complexes were consistent across five different detergents. However, it is unclear if, or to what extent, less harsh detergents such as NP-40 solubilize protein complexes associated with the PSD, as opposed to cytoplasmic proteins or those in the PSD periphery.

**Figure 1:**
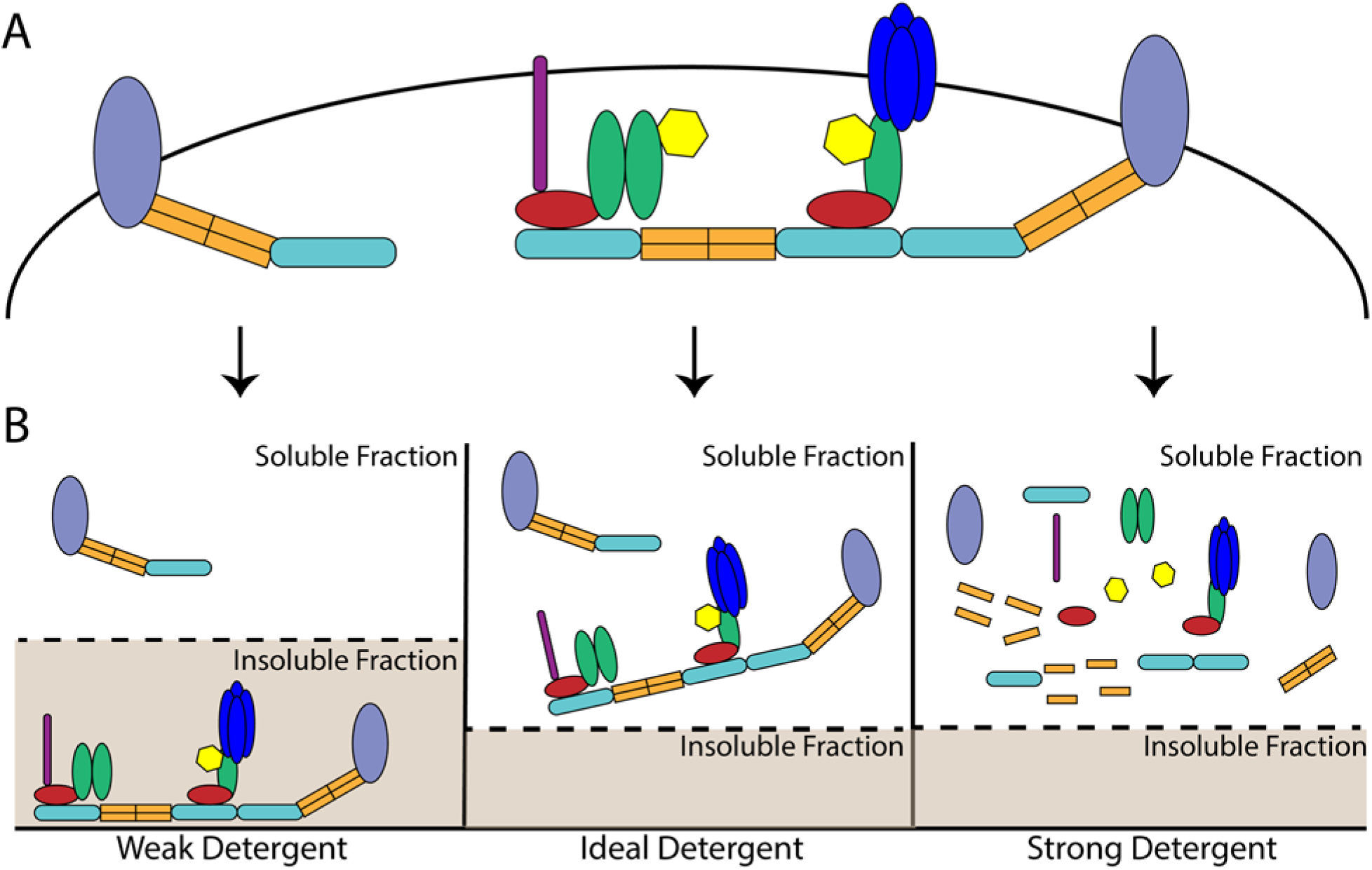
The impact of detergents on multiprotein complexes: (A) Organization of theoretical multiprotein complexes prior to solubilization. (B) The impact of distinct detergents on the observed abundance of multiprotein complexes. The ideal detergent will fully solubilize all proteins while also maintaining protein co-associations. By comparison, a weak detergent will maintain protein co-associations but solubilize only a portion of the proteins of interest while a detergent that is too strong will simultaneously solubilize all proteins and disrupt protein co-associations.

Here, we sought to quantify the ability of DOC, NP-40 and Triton to solubilize PSD-associated proteins while also maintaining protein complexes, to reconcile recent results using these detergents. We first confirm that synaptic protein complexes are only partially solubilized by NP-40 or Triton, but fully soluble in DOC. While almost all protein complexes are detected across all detergent conditions, many complexes are differentially abundant in distinct detergents. For example, complexes containing SynGAP are more abundant in DOC, while Homer_mGluR5 co-associations are detected to a much greater degree in NP-40 or Triton.

Importantly, activity-dependent changes in protein co-associations are consistent across detergent types, suggesting that they reflect synaptic biology. However, the detection of these activity-induced changes is more or less feasible in each detergent based upon the baseline abundance of each protein co-association. We suggest a model in which liquid-liquid phase separated PSD proteins are partially solubilized by NP-40 or Triton, allowing for detection of high MW complexes. Moreover, protocols using NP-40 or Triton can result in detection of a greater number of activity-dependent interactions, and may prove beneficial for future studies.

## Results

### The impact of detergent selection on synaptic protein solubilization

To determine the ability of different detergents to solubilize PSD-associated proteins, we made identical synaptosome preps (30-32) by dividing the S1 phase into 3 equal aliquots (Figure 2A). The resulting P2 pellets were solubilized in distinct lysis buffers containing 1% of either DOC, NP-40, or Triton; the insoluble portion of each lysate was subsequently isolated by centrifugation and solubilized in an equal volume of sample buffer containing SDS+BME. The relative amount of 6 synaptic proteins in the lysate vs. pellet solubilized in equal volumes of buffer was then quantified by western immunoblot (Figure 2A). As expected, DOC completely solubilized all PSD-associated proteins measured (Figure 2B). In contrast, NP-40 or Triton solubilized only a portion of the same synaptic proteins. Moreover, the relative amount of protein solubilized by these detergents varied based upon the protein of interest; NP-40 and Triton solubilized almost 90% of total mGluR5, but only 30-45% of GluR1, Homer1, NMDAR1, and PSD-95 and only 15% of SynGAP (Fig 2C-H). Collectively, these results demonstrate that all three detergents can solubilize synaptic proteins, albeit to varying degrees.

**Figure 2:**
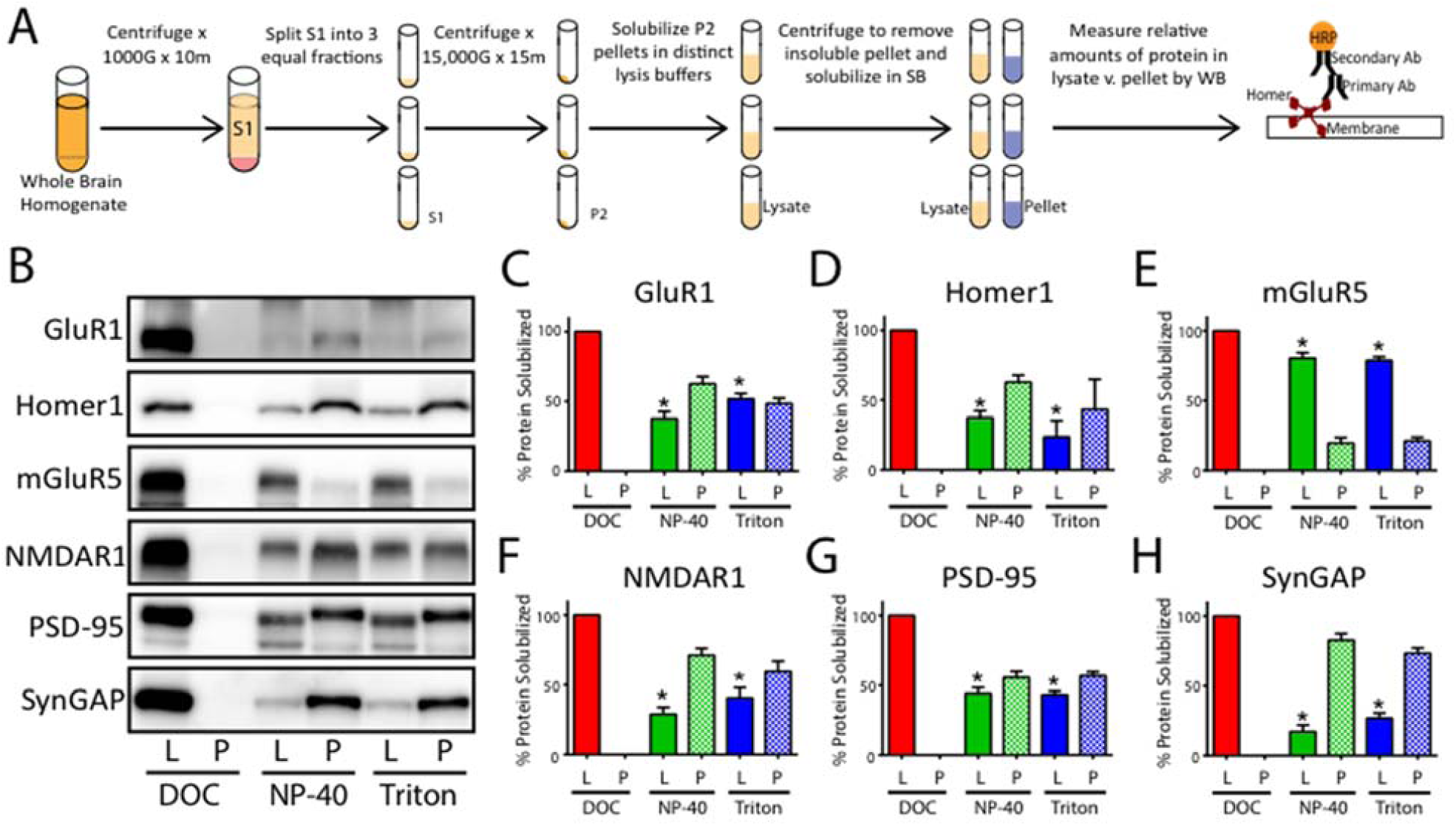
The impact of detergent selection on synaptic protein solubilization: (A) Workflow.(B) Representative western blots comparing the relative amount of 6 synaptic proteins in the soluble lysate (L) and insoluble pellet (P). (C) Quantification of N=3 blots including those shown in B. *p <0.05 relative to DOC lysate; Student’s t-test.

### The impact of detergent selection on synaptic protein interactions

While DOC (pH 9.0) best solubilized synaptic proteins, it may simultaneously disrupt protein co-associations. To examine the effect of detergent, as well as other commonly reported lysis buffer reagents (Tris vs. HEPES buffer and the use of calcium chelators) on protein co-associations, P2 preps were made from the same starting material and subsequently solubilized in 10 unique lysis buffers. The relative amount of selected protein co-associations was then quantified by immunoprecipitation detected by flow cytometry (((IP-FCM), Figure 3A)(33, 34)).

**Figure 3:**
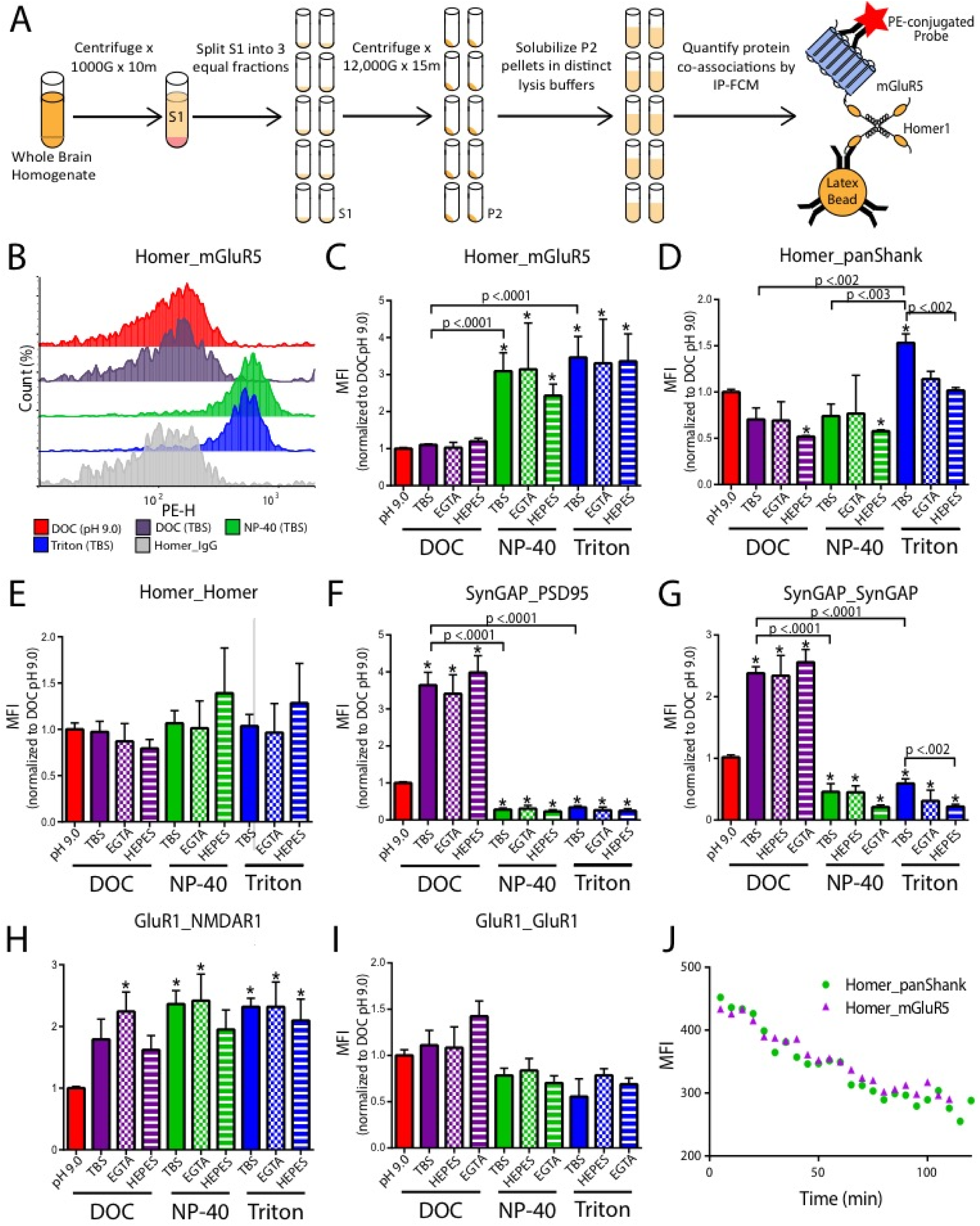
The impact of detergent selection on synaptic protein interactions. (A) Workflow. (B) Representative histograms of Homer_mGluR5; IP-FCM data are reported in the form “IP antibody_Probe antibody”. (C). Quantification of three IP-FCM experiments including those shown in B. (D-I) Quantification of IP-FCM experiments for N=3 experiments. (J) Quantification of Homer-containing complexes from reserve lysate that was allowed to warm to room temperature over the course two hours. Note decreased MFI values as time progresses indicating dissociation of Homer complexes when lysate is not maintained at 4 °C.*p <0.05, paired Student’s t-test.

We initially examined the effect of lysis buffer reagents on Homer_mGluR5, a well characterized synaptic protein interaction known to play an important role in glutamatergic signaling (19, 26, 29, 35). Significantly more Homer_mGluR5 co-association was detected in NP-40 and Triton compared to DOC (Figure 3B, C). Moreover, the fluorescent intensity histogram for Homer_mGluR5 in DOC conditions overlapped with that of the Homer_IgG control, indicating that that this interaction was at the lower limit of detection in DOC (Figure 3B). Homer_panShank was also significantly higher in Triton-containing lysis buffers compared to DOC or NP-40 (Figure 3D), whereas Homer_Homer, was not significantly affected by detergent selection, buffer choice, or the presence of calcium chelators (Figure 3E). Conversely, SynGAP containing interactions exhibited significantly higher median fluorescent intensities (MFIs) in DOC (Figure 3F-G). Interestingly, the use of DOC at a lower pH (7.4) led to even stronger detection of SynGAP interactions. For NP-40 or Triton containing lysis buffers, the use of HEPES resulted in lower MFIs, suggesting that these detergent/buffer combinations are the least appropriate for measuring SynGAP containing interactions. The use of DOC (pH 9.0), however, led to decreased detection of GluR1_NMDAR1 co-association when compared to all other conditions (Figure 3H). GluR1_GluR1 was detected at slightly higher level in DOC containing lysis buffers, though this result was not statistically significant (Figure 3I).

While DOC at a lower pH (7.4) was more effective at detecting several interactions than DOC at a high pH (9.0), this lysis buffer would solidify at 4°C if left without agitation (but maintained fluidity at room temperature). While seemingly intuitive that it is necessary to keep lysate at cold temperatures to maintain protein co-associations, we wanted to confirm that this was true. To this end, a P2 prep was solubilized in NP-40 and the amount of Homer_mGluR5 and Homer_panShank co-association was quantified by IP-FCM every minute for two hours.

The average MFI decreased by almost 50% (Figure 3J). This result demonstrates that it is necessary to maintain lysate at cold temperatures, making the use of DOC at room temperature unsuitable for protein co-association studies. Moreover, as EGTA did not significantly affect protein co-associations and the use of HEPES resulted in lower MFIs, the experiments below focus on lysis buffers containing DOC (pH 9.0), NP-40 (TBS), and Triton (TBS).

### Size exclusion chromatography reveals more abundant high MW complexes in DOC

P2 preps were made from the same starting biomaterial and multiprotein complexes were separated by size using Size Exclusion Chromatography (SEC). The relative amounts of synaptic proteins in each fraction were then detected by western immunoblot (Figure 4A). High MW complexes (i.e. fractions 1-4) containing Homer, mGluR5, PSD-95, and SynGAP were present in all detergent conditions. Similar to previous experiments, NP-40 and Triton yielded largely similar results (Figure 4B,C). DOC, however, yielded a distinct SEC profile characterized by a greater percentage of each protein in earlier fractions, and a shift toward higher apparent molecular weights of the low-MW peaks for Homer, mGluR5 and SynGAP. This result indicates that all detergents solubilized at least some large MW complexes. Counterintuitively, DOC solubilized more high MW complexes and complexes composed of the same proteins exhibited higher apparent MWs in DOC when compared to NP-40 or Triton.

**Figure 4:**
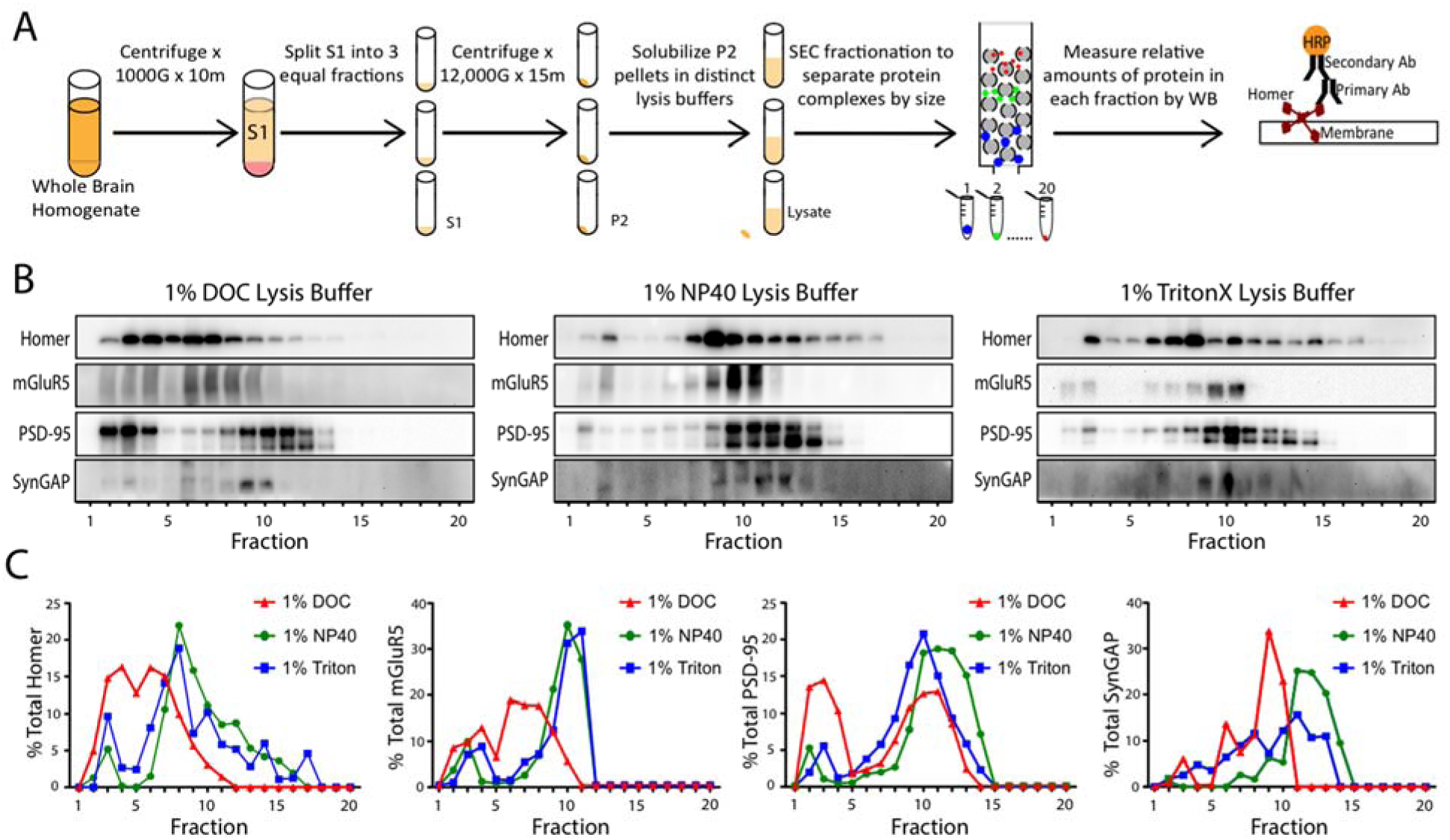
Detergent affects SEC profiles of synaptic proteins. (A) Workflow. (B) Representative western blots of synaptic proteins solubilized in different detergents and separated by SEC fractionation. (C) Quantification of N=2 blots, including those shown in B. Note, that while DOC lysis buffer produced more proteins in high MW fractions (i.e. 1-8), NP-40 and Triton conditions also had protein in high MW fraction.

### IP-FCM detects protein co-associations from high MW fractions

To determine if the co-associations observed in previous experiments are found in large MW complexes, SEC fractionation was performed on P2 preps solubilized in DOC, NP-40, or Triton. The amount of co-association in pooled fractions was then quantified by IP-FCM (Figure 5A). Consistent with previous results, Homer_mGluR5 co-association was only observed in NP-40 or Triton conditions (Fig 5B-E). Critically, the majority of this co-association was detected in fractions 1-4 and 5-8 (Figure 5E), despite the majority of mGluR5 being observed in fractions 9-12 by SEC-western (Figure 4C). This fact that the IP-FCM signal for fractions 1-4 was equivalent to (NP-40) or higher than (Triton) the signal observed in the lysate is remarkable considering that only a small minority of total protein was present in these fractions by SEC-western (Figure 4B).

**Figure 5:**
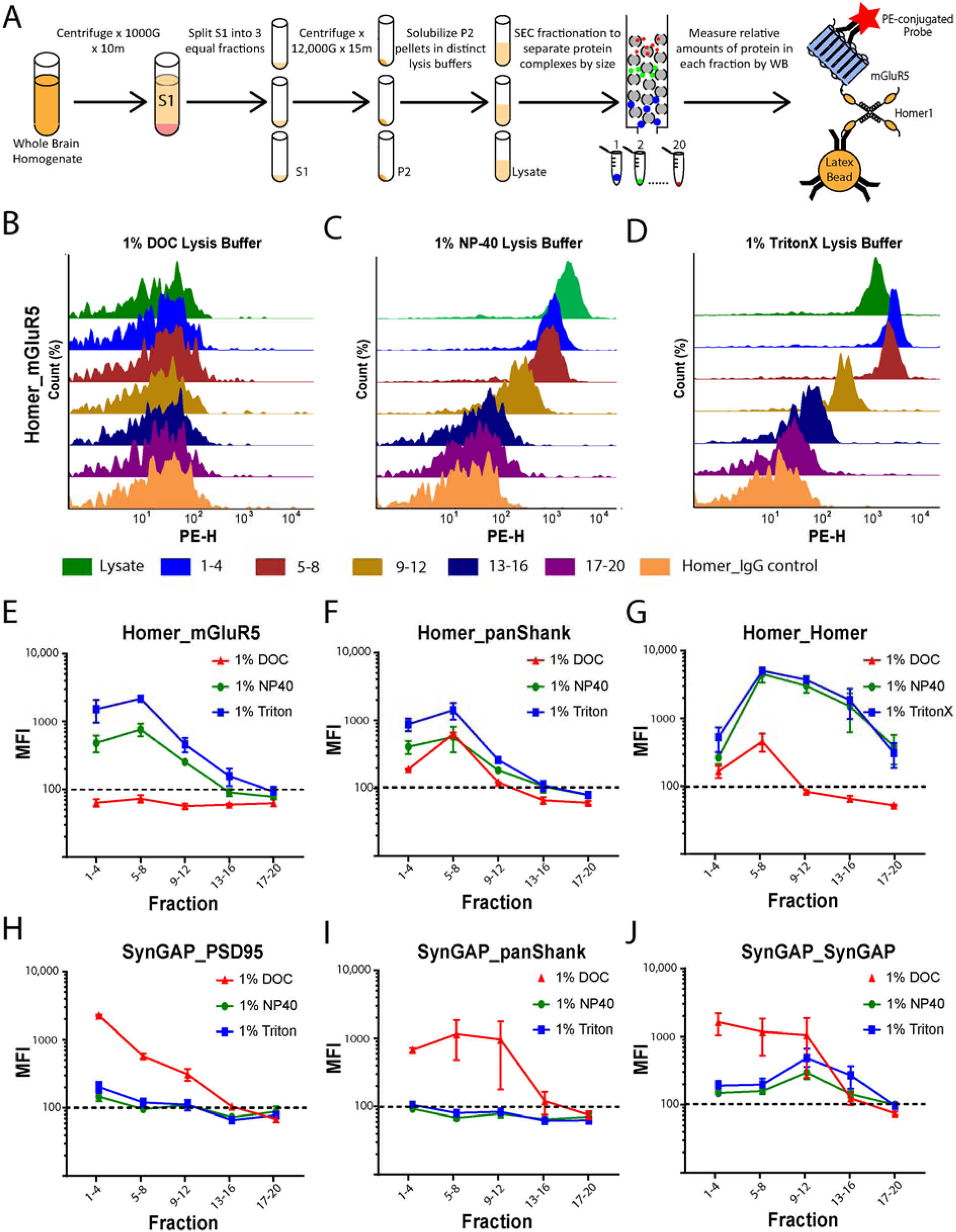
Co-associated proteins are found in large MW complexes. (A). Workflow. (B-D). Representative histograms showing the observed abundance of Homer_mGluR5 co-association across 5 groups of pooled fractions as well as unfractionated lysate controls. (E-J) Quantification 6 protein co-associations, including data shown in B. N=2-3.

Moreover, Homer_Homer detection by IP-FCM largely mirrored the SEC-western results, with signal spread over all fractions for both NP-40 and Triton, peaking in fraction 8. These results clearly show that Homer_mGluR5 co-association detected by IP-FCM is derived from high-MW SEC fractions, despite only a small minority of total protein residing there.

We performed parallel analysis of several other co-associations. Homer_panShank co-association was detected primarily in high MW fractions across all three detergent conditions (Figure 5F). SynGAP_PSD95 co-association (Figure 5H) was most abundant in the highest MW fractions (1-4) in all detergent conditions, and was detectable in fractions 1-4 in NP40 and Triton despite very little high-MW SynGAP by SEC-western (Fig 4B). While the MFI value for this co-association was markedly higher in DOC, consistent with the increased solubility reported above (Figure 2H), the signal in NP40 and Triton was significantly above background (IgG controls, not shown). Conversely, SynGAP_panShank co-association was detected solely in DOC conditions. The MFI values for this co-association were relatively equal in fractions 1-12, suggesting a range of high and medium MW complexes. SynGAP_SynGAP was detected in all three detergent conditions across a wide range of fractions, with again, the highest MFI values occurring in DOC conditions. Collectively, these data demonstrate that DOC, NP-40, and Triton solubilize and maintain high MW multiprotein complexes, though the use of DOC vs. NP-40/Triton results in differential abundance of specific complexes.

### Activity-dependent changes in protein co-associations, when detectable, are similar in all detergent conditions

As the use of DOC vs. NP40/Triton clearly alters the abundance of specific protein co-associations, it is plausible that detergent selection also results in distinct experimental outcomes when measuring activity-dependent changes in synaptic protein co-associations. For example, recruitment of proteins into the Triton insoluble portion of the PSD might be observed as *decreased* co-association in Triton/NP40 lysates but *increased* co-association in DOC lysates (a ‘source-sink’ model). Alternatively, a more labile protein co-association (such as Homer_mGluR5) may be disrupted by DOC at baseline, such that activity-dependent changes would not be detectable at all. To determine the effect of detergent selection on our ability to measure activity-dependent changes in synaptic protein co-associations, we prepared hemisected cortical slices from p20-30 CD1 mice and stimulated them with either KCL or aCSF control for 5 minutes. Following homogenization, the S1 phase was divided into three, and the resulting P2 pellets were homogenized in lysis buffers containing either 1% DOC, NP-40 or Triton. Selected, known-activity-dependent changes in protein co-associations were then quantified by IP-FCM (Figure 6A). Following stimulation, we observed consistent dissociation of protein interactions across all three conditions, though certain dissociations were better detected in specific detergents (Figure 6B). For example, Homer_mGluR5 dissociation was detected in all conditions, though to a greater magnitude in NP-40 or Triton, since at baseline in DOC, this interaction was almost at a floor (background MFI≈100). Similarly, dissociation of SynGAP_PSD95, SynGAP_panShank, and SynGAP_SynGAP was observed in all three conditions, but to the greatest extent in DOC. For almost all interactions (except Homer_Homer), we observed the greatest magnitude of change in protein co-association in the detergent with the highest baseline MFI. This result is logical, as here we are only measuring activity-dependent *dissociations*, and interactions with a lower baseline MFI have less room to decrease before hitting our lower limit of detection (∼100MFI).

**Figure 6:**
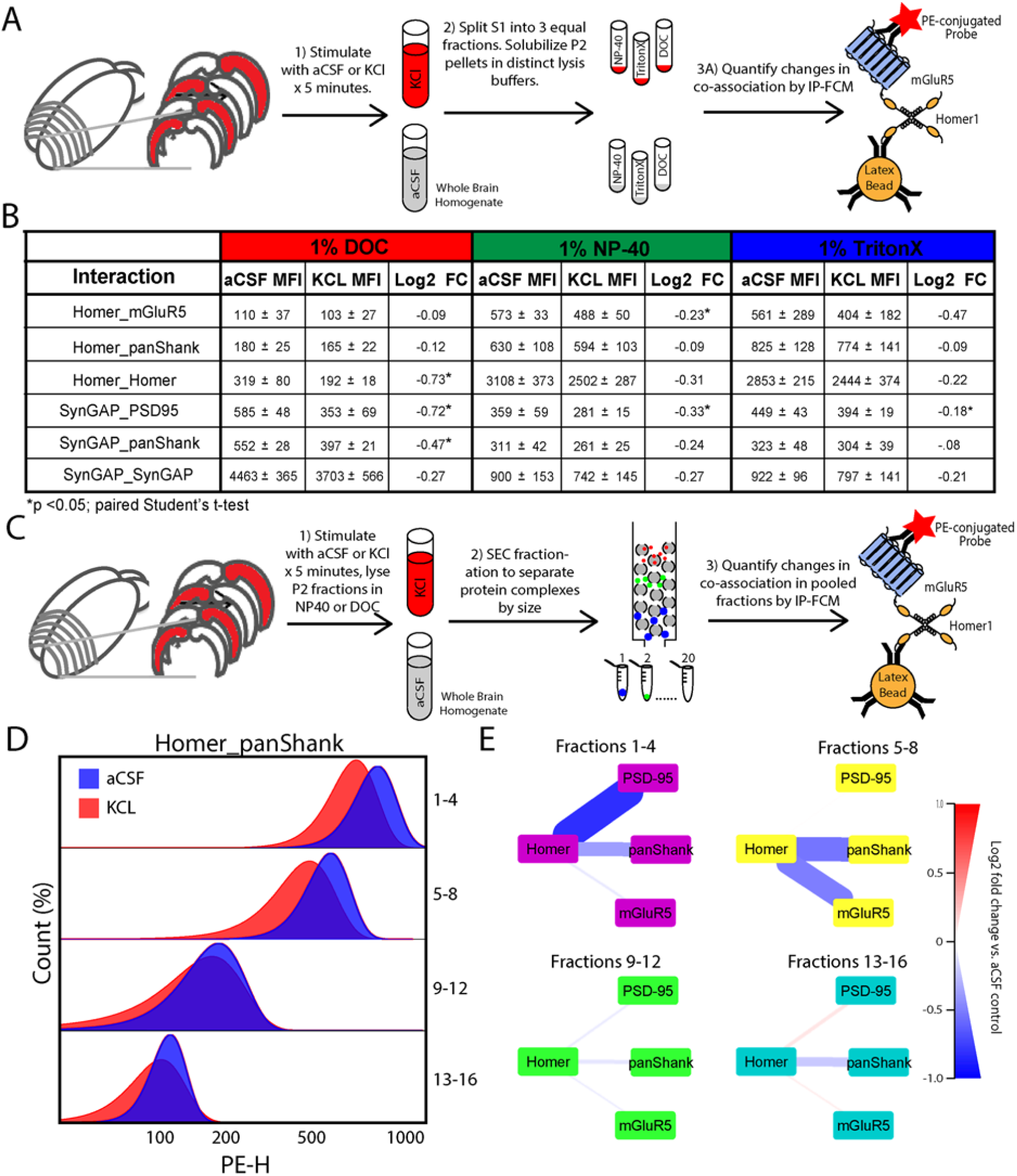
Activity-induce changes are similar across detergents and occur in high MW complexes. (A). Workflow for B. (B) Table showing MFI and fold change of each interaction measured. Note that while the MFIs at baseline (in aCSF) were different, changes in aCSF vs KCl were consistently of similar magnitude and direction. (C) Workflow for D and E. (D) Representative histograms show bead distributions for stimulated (KCL, red) and unstimulated (aCSF, blue) fractions in each pooled fraction. Note that within each pooled fraction, KCL stimulation elicited decreased association of Homer_panShank, with the largest dissociations occurring in high MW fractions (i.e. 1-4 or 5-8). (E) Node-edge diagrams showing Log2 FC for four Homer interactions in NP-40 lysis buffer (interactions were not detectable in DOC). The thickness and color of lines connecting protein nodes indicate the magnitude and direction of the fold change, with decreased association being colored blue, and increased association being red. N=2-3.*p <0.05, paired Student’s t-test.

To determine if the activity-dependent changes observed were occurring in large MW complexes, we prepared cortical slices from CD1 mice, stimulated with either KCL or aCSF, and homogenized the P2 pellets in lysis buffer containing either 1% DOC or NP-40. SEC fractionation was then performed on lysates and the relative amount of selected Homer interactions were quantified by multiplexed IP-FCM (Figure 6C). Following stimulation, we observed dissociation of Homer from mGluR5, panShank, and PSD-95 in NP-40 conditions.

Moreover, the majority of dissociations observed were detected in fractions 1-4 and 5-8 (Figure 6D, E), demonstrating that the activity-dependent changes detected in NP-40 conditions occur in large multiprotein complexes. The baseline MFI value for Homer interactions was markedly lower in DOC conditions, preventing detection of activity-dependent changes. Collectively, these results demonstrate that it is possible to measure activity-dependent changes in synaptic protein interactions in NP-40, Triton, or DOC, but the optimal detergent to use varies based upon the protein of interest.

### QMI reports widespread protein interaction network differences based on detergent

To better understand the effect of detergent on synaptic protein interaction networks, hemisected cortical slices from P20-30 mice were prepared and stimulated with either NMDA/Glycine (100/10 uM) for 5 min, a treatment previously demonstrated to produce widespread dissociation of synaptic protein complexes (28), or aCSF control. P2 preps from the same starting biomaterial were solubilized in lysis buffer containing either 1% DOC, NP-40, or Triton, and network level changes among an 18-member targeted synaptic protein interaction network were quantified by quantitative multiplex co-immunoprecipitation (QMI), a novel proteomics technique that allows for the simultaneous and quantitative measurement of the amount of co-association between large numbers of proteins (18, 33)(Figure 7A).

**Figure 7:**
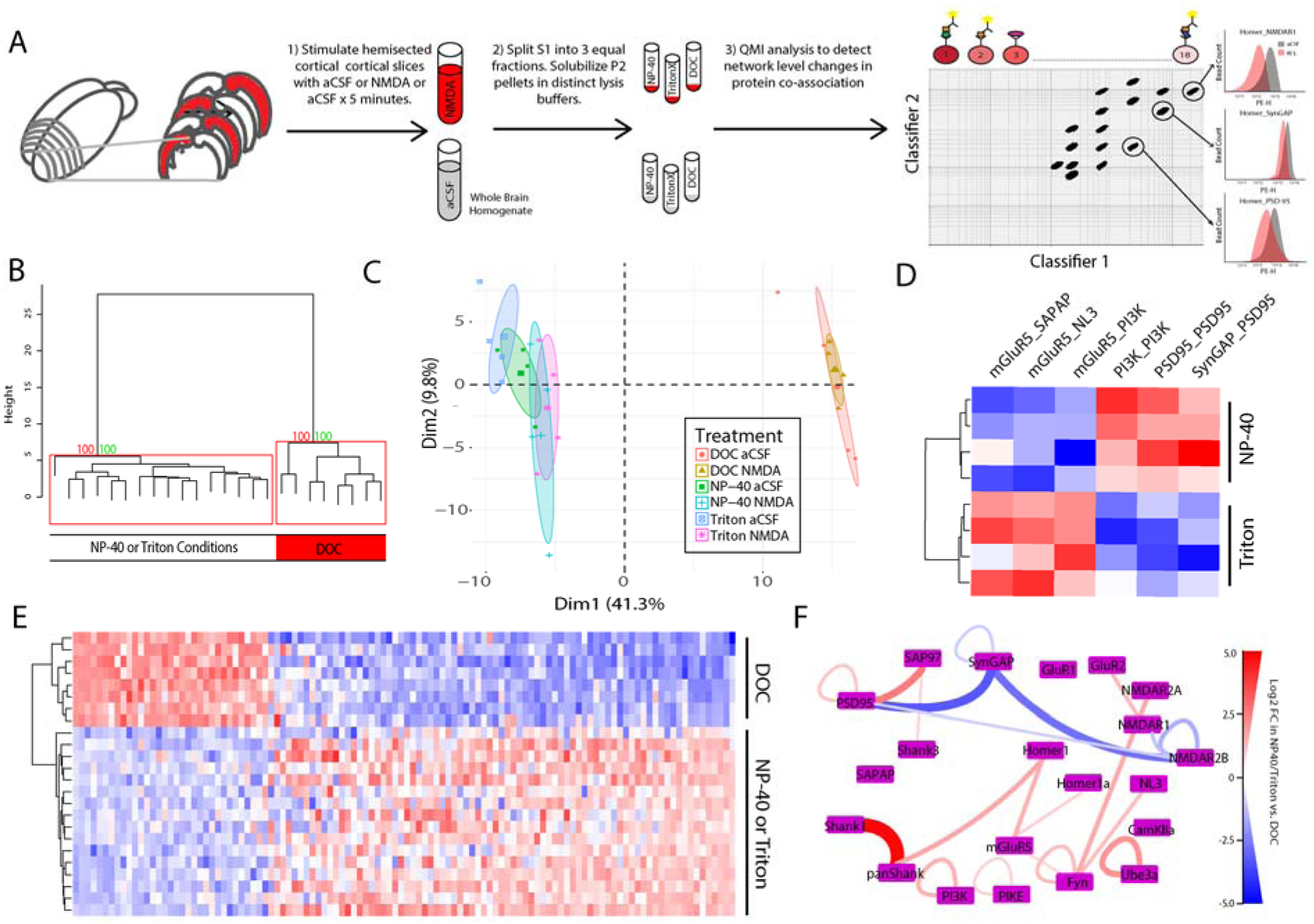
Multiplex IP-FCM detects more complexes in NP-40 or Triton Conditions: (A) Workflow. (B) Unsupervised hierarchal clustering of all conditions. Numbers at the branch points show the approximately unbiased (AU) p value calculated by multiscale bootstrap resampling; clusters with AU greater than 95 are boxed in red. (C) PCA of all conditions. Note DOC and NP-40/Triton conditions separated by both hierarchal clustering and PCA. (D) Heatmap of 6 interactions identified as significantly different between NP-40 and Triton conditions by ANC ∩ CNA analysis. (E) Heatmap of 112 interactions identified as significantly different between either DOC vs. NP-40 or DOC vs. Triton conditions. Note that hierarchal clustering again separated NP-40 and Triton conditions from DOC. (F) QMI map of the all interactions that exhibited greater than a >2 fold difference between between DOC and NP-40/Triton conditions. The thickness and color of lines connecting protein nodes indicate the direction and magnitude of the fold change with interactions that are higher in NP-40/Triton colored red and interactions that are higher in DOC colored blue.

Unsupervised hierarchal clustering of all detected interactions separated DOC conditions from NP-40 and Triton (Figure 6B). Similarly, PCA analysis revealed strong separation between DOC and NP-40/Triton along PC1, with little difference between NP40 and Triton (Figure 6C). By comparison, there was no clear distinction between stimulated and unstimulated conditions by hierarchal clustering, and only a small separation between stimulated and unstimulated conditions by PCA in NP-40 or Triton. To identify interactions that were significantly different between detergent conditions, we used two independent statistical approaches, adaptive non-parametric (ANC) analysis, and weighted correlation network analysis (CNA). Prior work with QMI data found that interactions identified independently by both analyses represent high confidence hits with low rates of false positives (18, 36). Comparing unstimulated NP-40 and Triton conditions, ANC∩CNA identified 6 interactions that were significantly different (Figure 6D). By comparison, ANC∩CNA identified 112 interactions that were significantly different between DOC and either NP-40 or Triton conditions (Figure 6E). For most protein co-associations, MFI values tended to be higher in NP-40 or Triton conditions, except for interactions involving SynGAP and NMDAR2B, which were markedly higher in DOC. To determine the interactions that most strongly differentiated DOC and NP-40/Triton conditions, we identified the interactions that averaged greater than a 2-fold change between conditions (Fig 6F). Shank1_panShank topped the list (MFI in NP-40 vs. DOC = 13,000 vs. 300), followed by SynGAP_PSD95 (300 vs. 4000) and SynGAP_NMDAR2B (100 vs. 600). Notably, Homer_mGluR5 and Homer_panShank were also detected at a higher level in NP-40/Triton, while NMDAR1_NMDAR2B was higher in DOC. Collectively, these data suggest that Homer_Shank complexes are better detected in NP-40/Triton, whereas SynGAP and NMDAR2B containing complexes can be more easily observed in DOC.

In order to better understand the effect detergent selection has on our ability to detect activity-induced changes in protein co-associations, we analyzed unstimulated vs. stimulated slices in each detergent separately. NMDA and aCSF conditions were separated by both hierarchal clustering and PCA in all detergent conditions (Figure S1), suggesting that we can detect network level changes in protein co-associations regardless of detergent selection. ANC∩CNA identified a total of 21 interactions that were significantly different following stimulation in at least one detergent condition. In DOC, we observed dissociation of 4 SynGAP-containing interactions (Figure 8A). Two of these dissociations, SynGAP_PSD95 and SynGAP_NMDAR1, were also observed in NP-40 and Triton conditions. ANC∩CNA identified 9 interactions that were significantly different following stimulation in NP-40 (Figure 8B). These changes were characterized by dissociation of Homer-, SynGAP-, and PSD-95-containing complexes, as previously reported (18). Similarly, we observed 12 significant changes following stimulation in Triton that were characterized by dissociation of Homer-, SynGAP-, and mGluR5-containing complexes (Figure 8C). To directly compare stimulation results in different detergents, the log2 fold change for every protein interaction significant in either condition was plotted in X-Y coordinates. Activity-dependent changes in NP-40 and Triton were largely similar (Figure 8D). By comparison, the activity-dependent changes observed in DOC vs. NP-40 were different (Figure 8E), with the fold change for 7 out of the 9 interactions that were significantly different in NP-40 falling below the level of detection in DOC (10% change, gray boxes).

**Figure 8:**
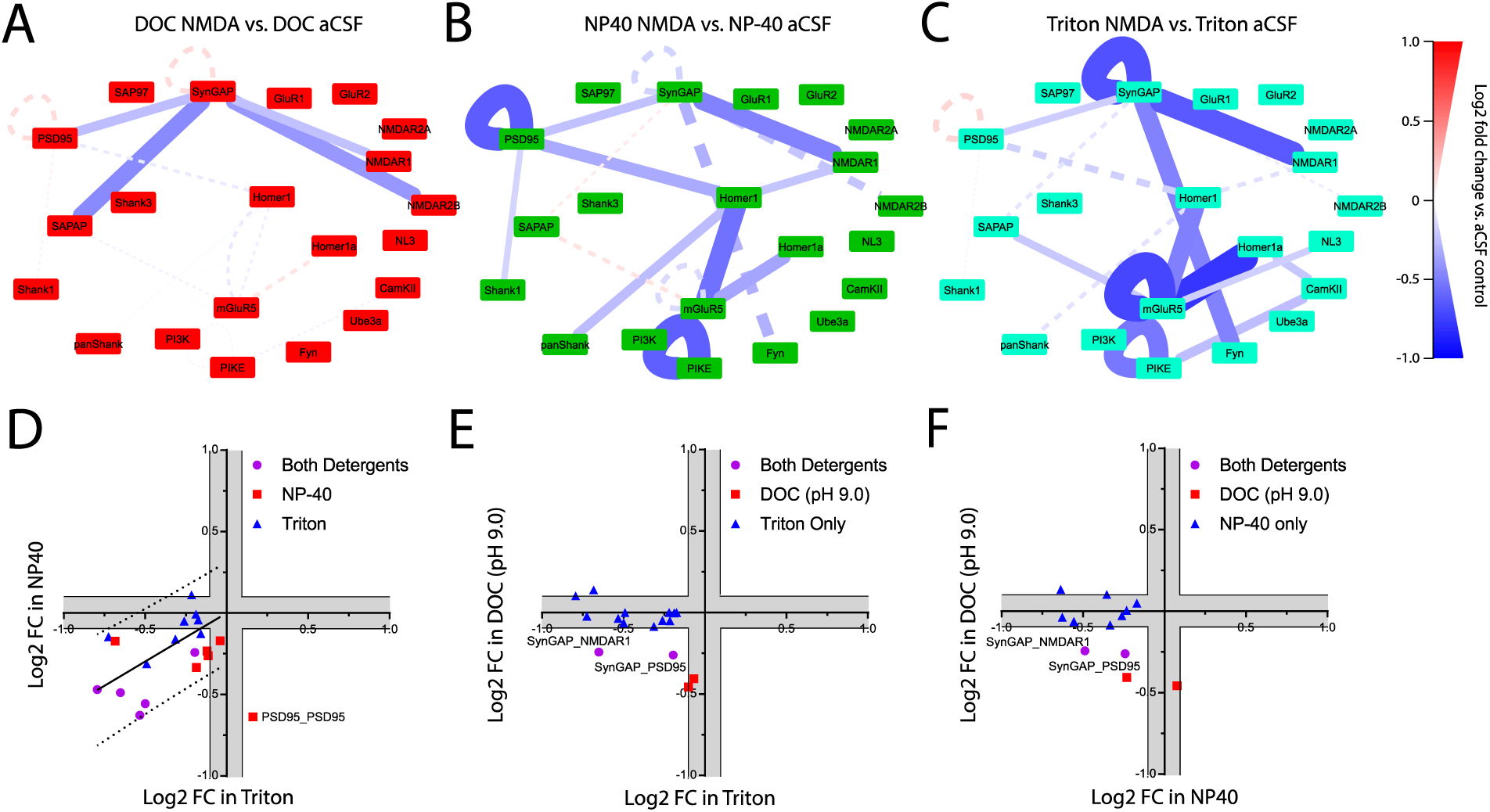
Activity-dependent changes in synaptic protein complexes across detergent conditions: To identify interactions that changed significantly following NMDA stimulation, each detergent condition was analyzed separately. Node-edge diagrams show all interactions identified as significantly different by ANC ∩ CNA analysis in 1% (A) DOC, (B) NP-40, or (C) Triton. The solid lines on each QMI map represent interactions that were significantly different in that specific detergent condition; dotted lines show the non-significant fold change for interactions that were significantly different in other detergent conditions to highlight the lack trends in the opposite direction. The color and thickness of lines connecting protein nodes indicate the direction and magnitude of the change. (D-F). To directly compare the observed activity-dependent changes in each condition, we plotted the Log2 fold change for every interaction significant under either condition. Gray areas represent less than a 10% change, the lower limit of detection for QMI.

Similarly, the fold change for 10 of the 12 interactions identified as significant in Triton fell below the level of detection (10% change, gray boxes) in DOC conditions (Figure 8F). Critically, we never saw an interaction that was strongly decreased in NP-40/Triton conditions, but strongly increased in DOC, which leads us to reject the ‘source-sink’ hypothesis of detergent pools and conclude that the same changes in protein co-associations are observed, regardless of detergent used.

Collectively, these results demonstrate that detergent selection strongly affects our ability to measure network level changes in the composition of multiprotein complexes at the glutamate synapse. The use of DOC results in greater detection of activity-dependent changes in SynGAP and NMDAR2B containing complexes. By comparison, activity-dependent changes in Homer and Shank complexes are exclusively detected in NP-40 or Triton. These data suggest that the use of less harsh detergents may enable the detection of novel, activity-dependent interactions.

## Discussion

In this study, we characterized the effect of DOC, NP-40, and Triton on the solubility and composition of synaptic multiprotein complexes. We also demonstrated how the use of different detergents can affect our ability to measure activity-dependent changes in protein co-associations. To accomplish this, we adopted a methodology of making multiple P2 preps from the same starting material by separating the S1 fraction into multiple, equal aliquots. This approach eliminates any variability that may that may exist between samples, and effectively isolates detergent as the experimental variable. Theoretically, the optimal detergent would both fully solubilize the PSD and also maintain protein co-associations (Figure 1). Here, we found that DOC fully solubilizes the PSD but also disrupts protein co-associations. By comparison, NP-40 and Triton only partially solubilized PSD associated proteins, but maintained certain interactions better than DOC.

These data beg the question, are the protein co-associations observed in NP-40 and Triton conditions localized to the PSD *in vivo* It is possible that the solubilized proteins in the NP-40 or Triton conditions represent a pool of peri-synaptic PSD-associated proteins that do not localize to the PSD. We find this unlikely for three reasons: first, we confirmed by SEC fractionation that both NP-40 and Triton solubilize high MW complexes that contain 4 well known PSD associated proteins: Homer, mGluR5, PSD-95, and SynGAP. Second, we recapitulated two well-known activity dependent interactions that have been localized to within the PSD, dissociation of Homer_mGluR5 (29) and SynGAP_PSD95 (22). We find it unlikely that the high MW multiprotein complexes observed by SEC-western in NP-40 or Triton conditions are both composed of PSD-associated proteins, and exhibit activity-dependent dynamics similar to the PSD, but are not actually part of the PSD. Finally, given that NP40/Triton does not solubilize a large portion of total synaptic protein, we considered the possibility that the *dissociation* of proteins previously reported (18) in NP40/Triton could reflect a depletion of proteins from a hypothetical “NP40/Triton-solubilized perisynaptic pool”, and recruitment of proteins into the PSD. In this “source-sink” model, we would expect to see increases in co-associations in DOC solubilized complexes, reflecting recruitment of perisynaptic proteins into the PSD. Instead, we observed similar changes (i.e. in the same direction) in protein complex abundance in all detergent conditions, although detection of activity-dependent changes was more or less feasible based on the baseline MFI of interaction in each detergent. We therefore posit that NP-40 and Triton can solubilize a portion, albeit not all, of the PSD, and the changes in synaptic protein complexes we observe are directly relevant to PSD biology.

Recent evidence suggests that the PSD is a liquid-liquid phase separated (LLPS) structure, in which weakly interacting proteins spontaneously segregate to form a discrete subcellular compartment, lacking a membrane (14, 37). In this model, proteins are consistently diffusing between the dense LLPS structure and the less concentrated aqueous surrounding. We propose a model in which NP-40 and Triton detergents permit diffusion of synaptic protein complexes from a core LLPS structure into the aqueous surroundings; proteins such as SynGAP that bind with multiple components of the PSD and are tightly integrated into the LLPS structure diffuse to a lesser extent, while more peripheral proteins such as mGluR5 with single binding sites anchoring them diffuse more rapidly, leading to a greater level of solubilization. Future work testing this hypothesis with an in vitro reconstructed system (38) could clarify the origin of the multiprotein complexes observed here.

In regards to the best detergent for future studies, our data demonstrate that most interactions are detected at a higher level in NP-40 or Triton, whereas SynGAP-and NMDAR2B-containing interactions were much more abundant in DOC. Most notably, Homer_mGluR5, Homer_panShank, and Shank1_panShank were all markedly lower in DOC. It is unclear why these proteins would co-migrate in high MW fractions in DOC (Figure 4B), but not be detected physically interacting by IP-FCM (Figure 5B). It is possible that the limitations of IP-FCM-specifically the possibility that the required antibody binding could be disrupted by high pH or occluded by steric interference in DOC buffers-may prevent us from measuring protein complexes that are, in fact present. However, for our purposes of identifying a lysis buffer most useful for biochemical experiments, we conclude that DOC limits our ability to measure Homer and Shank containing interactions.

When we did examine activity-dependent changes in different detergents, we observed widespread dissociation of Homer, Shank and SynGAP containing complexes. Only two interactions were detected sufficiently in all detergent conditions to reach the stringent ANC∩CNA significance criteria: dissociation of SynGAP_PSD95 and SynGAP_NMDAR1. The dispersion of SynGAP from the PSD following activity has been previously demonstrated by both electron microscopy (20) and co-immunoprecipitation (39), and this activity-dependent interaction is considered to be an important first step for induction of long term potentiation (21, 40). According to the “Slot hypothesis” (22), the rapid dissociation of SynGAP from PSD-95 frees the PDZ-binding domain of PSD-95, allowing for the additional recruitment and binding of AMPARs, critical to short-term potentiation (41). Indeed, previous work using QMI has demonstrated that longer stimulation elicits recruitment and stabilization of AMPA receptors(18), consistent with the slot hypothesis. In addition to SynGAP interactions, we also observed widespread dissociation of Homer, Shank, and mGluR5 complexes in NP-40 or Triton conditions. The co-clustering of NMDAR’s and mGluR5 through a PSD-95/Shank/Homer complex has been shown to alter the physiology of these receptors, allowing both to respond to lower activation thresholds (35, 42, 43). As this coordination is dependent on both Homer binding to mGluR5 (35) and PSD-95 binding to NMDAR2B(42), it is possible that widespread dissociation represents a homeostatic response, designed to maintain signal strength at an appropriate level. Alternatively, widespread dissociation of scaffold proteins may allow for the competitive capture of dendritic proteins, expansion of the synapse, and an increase in synaptic strength (22). Decreased association of Homer_mGluR5 itself has been crucially implicated in both normal mGluR5 signaling and the pathophysiology of Fragile X syndrome (19, 29, 44), further supporting the physiological relevance of our findings in NP-40 and Triton conditions.

Collectively, our data support a model in which strong synaptic stimulation elicits widespread dissociation of multiprotein complexes localized to the PSD. Moreover, many of the activity-dependent interactions observed involve the dissociation of Homer-containing complexes, suggesting that Homer1 plays an essential role in glutamatergic signaling. Our results also demonstrate that it is critical for researchers to acknowledge the limitations of each detergent and carefully consider how detergent selection may affect experimental outcomes. The use of non-traditional solubilization protocols may result in the identification of novel protein co-associations and a more accurate understanding of the molecular mechanisms underlying synaptic function.

## Methods

### Animals

CD-1 (RRID:IMSR_CRL:22) mice were originally obtained from The Jackson Laboratory (Bar Harbor, Maine) and maintained in an in-house breeding colony. All mice were separated by sex, and housed with littermates in thoren cages, with no more than five mice/cage. Food and water was provided ad libitum. For slice experiments, only p21-30 mice (both male and female) were used. To minimize suffering, mice were anesthetized with isoflurane and decapitated under deep anesthesia. The use and care of animals complied with the institutional guidelines of Office of Animal Care at the Seattle Children’s Research institute (protocol# 15580).

### Lysate preparation

A graphical representation of each experimental design is included in each figure. Briefly, for P2 preps, mice were deeply anesthetized with Isofluorane, brains were removed, and tissue was homogenized in 0.32M sucrose in 5 mM 4-(2-hydroxyethyl)-1-piperazineethanesulfonic acid (HEPES) buffer with protease/phosphatase inhibitor cocktails using 12 strokes of a glass-teflon homogenizer. The homogenate was immediately centrifuged x1,000G for 10 minutes at 4°C. The S1 was then divided into the appropriate number of aliquots, and spun at 12,000G for 15 minutes at 4°C. The resulting P2 pellet was then resuspended in the appropriate lysis buffer x 15 minutes (Supplemental Table 1). Lysate was then centrifuged at 12,000G for 15 minutes to remove insoluble portions and protein concentration in the supernatant was determined using a Pierce BCA kit (Pierce, 23225).

### Brain dissection, slice preparation, stimulation, and lysis

Mice were deeply anesthetized with Isofluorane, brains were removed, and coronal cortical slices were sectioned at 400 µm thickness using a vibratome. Slices were immediately hemisected with a sharp razor blade and each half placed in an alternate treatment group with treatment groups being arbitrarily assigned. Slices were initially recovered in NMDG-protective recovery solution (93 mM NMDG, 2.5 mM KCl, 1.2 mM NaH2PO4, 30 mM NaHCO3, 20 mM HEPES, 25 mM glucose, 2 mM thiourea, 5 mM Na-ascorbate, 3 mM Na-pyruvate, 0.5 mM CaCl2.4H2O, and 10 mM MgSO4.7H2O; titrated to pH 7.4 with concentrated hydrochloric acid) for 10–15 min at 32–34°C, then transferred to a modified HEPES holding solution [92 mM NaCl, 2.5 mM KCl, 1.2 mM NaH2PO4, 30 mM NaHCO3, 20 mM HEPES, 25 mM glucose, 2 mM thiourea, 5 mM Na-ascorbate, 3 mM Na-pyruvate, 2 mM CaCl2.4H2O, and 2 mM MgSO4.7H2O; pH 7.4] for an additional 60–90 min recovery at room temperature using the protective recovery method (45). For KCl stimulation, slices were incubated at 37°C in 50 mM KCl or control HEPES-aCSF for 5 min. Following stimulation, tissue was homogenized and processed as previously described.

### IP-FCM

IP-FCM was performed as described previously (33, 46). CML latex microspheres (Invitrogen #<u>C37255</u>, USA) were activated with EDAC (1-ethyl-3-(3-dimethylaminopropyl) carbodiimide HCl; Pierce, USA), and coupled to 50 µl of 0.5 mg/ml antibody for 3 hours at room temperature. Probe antibodies were biotinylated at 0.5 mg/ml with EZ-link Sulfo-NHS-Biotin (Thermo, USA). Following solubilization of P2 pellets in distinct lysis buffers, protein concentrations were normalized by BCA assay and 2.5e^4^ antibody-conjugated beads were added to the lysate and incubated overnight at 4 C with rotation. The following day, beads were washed (x3) in Fly-P buffer (50 mM Tris (pH 7.4), 100 mM NaCl, 1% bovine serum albumin, and 0.02% sodium azide), biotinylated probe antibodies were added for 2h on ice, followed by washing (x3), and incubation with 1:200 streptavidin-phycoerythrin (PE, Biolegend 405204) for 30m. CML beads were then analyzed on a flow cytometer (Novocyte). MFI values and bead distributions were used for analysis.

### Size exclusion chromatography

Following solubilization in distinct lysis buffers, lysate protein concentrations were normalized by BCA. Lysates were then injected in a Superose 6 Increase 10/300 GL) with flow rate of 1 mL/min in the appropriate lysis buffer. Fractions were advanced at 1.5-min intervals. While absorption was monitored, the presence of 1% detergent in the flow through increased absorption making detection of proteins in the effluent impossible. For western blot analysis, proteins in each fraction were then concentrated using methanol precipitation. For detection of protein co-associations, 4 samples were pooled for IP-FCM analysis.

### QMI

QMI experiments were performed as described previously ((18, 28, 36)); all procedures were performed at 4°C or on ice. Briefly, a master mix containing equal numbers of each antibody-coupled Luminex bead was prepared and distributed into post-nuclear cell lysate samples, in duplicate. Protein complexes were IP’d from samples containing equal amounts of protein overnight at 4°C, washed twice in ice-cold Fly-P buffer and distributed into twice as many wells of a 96-well plate as there were probes, on ice. Biotinylated detection antibodies were added and incubated for 1 h, with gentle agitation at 500 rpm in a cold room (4°C). For antibody panel details, see (18). Following incubation, microbeads and captured complexes were washed three times in Fly-P buffer using a Bio-Plex Pro II magnetic plate washer at 4°C. Microbeads were then incubated for 30 min with streptavidin-PE on ice, washed three times, and resuspended in 125 μL of ice-cold Fly-P buffer. Fluorescence data were acquired on a customized, refrigerated Bio-Plex 200 using Bio-plex Manager software (version 6.1). The instrument was calibrated and routinely validated according to the manufacturer’s recommendations. Data files were exported in both Microsoft Excel and XML formats for further processing. Each experiment generated a 17 × 20 matrix of capture and detection antibodies, measuring 378 protein co-associations.

### Data preprocessing and inclusion criteria

For each well from a data acquisition plate, data were processed to (i) eliminate doublets on the basis of the doublet discriminator intensity (> 5000 and < 25 000 arbitrary units; Bio-Plex 200), (ii) identify specific bead classes within the bead regions used, and (iii) pair individual bead PE fluorescence measurements with their corresponding bead regions. This processing generated a distribution of fluorescence intensity values for each pairwise measurement. XML output files were parsed to acquire the raw data for use in MATLAB. No specific analysis was performed on the data to test for outliers.

### ANC

Adaptive non-parametric analysis with empirical alpha cutoff (ANC) is used to identify high-confidence, statistically significant differences (corrected for multiple comparisons) in bead distributions on an individual protein interaction basis. ANC analysis was conducted in MATLAB (version 2013a) as described in (36). As previously reported, we required that hits must be present in > 70% of experiments (typically three out of four) at an adjusted *p* < 0.05. The α-cutoff value required per experiment to determine statistical significance was calculated to maintain an overall type I error of 0.05 (adjusted for multiple comparisons with Bonferroni correction), with further empirical adjustments to account for technical variation (36). No assessment of normality was carried out as ANC analysis is a non-parametric test.

### Weighted correlation network analysis

Weighted correlation network analysis (CNA) is a second, independent statistical method to identify co-varying modules of protein interactions, and to then correlate those modules to experimental variables. Importantly, CNA relies on different assumptions than ANC, querying not what an individual interaction is doing, but what coordinated sets of interactions do together, as a unit. Remarkably, the two different approaches converge on a set of ‘high-confidence’ protein interactions. Bead distributions used in ANC were collapsed into a single median fluorescent intensity (MFI), which was averaged across technical replicates and input into the weighted gene correlation network analysis package for R studio (version 3.4.1)(47, 48). Data were filtered to remove weakly detected interactions (‘noise’, MFI <100), and batch effects were removed using the COMBAT function for R (48) with ‘experiment number’ as the ‘batch’ input. Post-Combat data were log_2_ transformed prior to CNA analysis. Soft thresholding using a power adjacency function was used to determine the power value resulting in the best approximation of scale-free topology, and the minimum module size was set to between 5 and 10, with the goal of generating a manageable number of modules (generally between 5 and 10). Protein interactions whose behavior was tightly correlated across experiments were assigned to arbitrary color-named modules by the weighted gene correlation network program. Modules whose eigenvectors significantly (*p* < 0.05) correlated with a given experimental trait (e.g., ‘KCl stimulation’, coded 1 vs. control, coded 0) were considered significantly correlated with the specific trait, and protein interactions belonging to modules of interest were defined as those with a probability of module membership (p.MM) < 0.05, as defined in the weighted gene correlation network program (47).

### ANC ∩ CAN

To ensure reporting of a core, high-confidence group of protein interactions, ANC data and CNA data were merged as described previously (36) to produce a high-confidence set of interactions that were both individually, statistically significantly different in comparisons between experimental groups, and that belonged to a larger module of co-regulated interactions that was significantly correlated with experimental group.

### Clustering and principal component analysis

For hierarchical clustering, log_2_ transformed post-combat data were clustered using the Ward’s method with a Euclidean distance matrix using the statistical package flashClust in R studio. Principal component analysis was performed on the same data in R studio using the prcomp function.

### Data visualization

QMI maps were generated to visualize only ANC ∩ CNA merged significant hits for all experiments using the open network resource Cytoscape (version 3.6.1). Mean fold changes, calculated from replicate experiments with significant fold change differences (by ANC analysis), were used to generate diagram edges. For protein pairs with significant changes in multiple measurements with different epitope combinations, the measurement with the greatest absolute log_2_ fold change value was selected for visualization (36). For heatmap visualization of fold change data, Heatmap.2 in R was used. For heatmap visualization of MFI values of CNA module members, log2 transformed data were input into the Heatmap program in R studio, which normalized the data by row for visualization of multiple analytes spanning a 3-log range.

### Western Immunoblot

For all western blot experiments, gels were transferred onto polyvinylidene difluoride (Millipore) membranes, blocked in 4% milk or bovine serum albumin in TBST (0.05 M Tris, 0.15 M NaCl, pH7.2, 0.1% (v/v) Tween20) for 60 min at room temperature, and primary antibodies were applied overnight at 4°C in blocking medium. Primary antibodies for western blots were diluted as follows: GluR1 (1504 poly, EMD Millipore, 1:1000), Homer1 (AT1F3, LS Bio, 1:1000), mGluR5 (ab5675, Millipore,1:1000), NMDAR1 (MAB363, Millipore,1:1000), PSD-95 (K28/43, NeuroMab, 1: 1000), SynGAP (D20C7, Cell Signaling, 1:1000) After washing and probing with the appropriate species-specific secondary horseradish peroxidase conjugated antibodies, blots were imaged using Pierce Femto detection reagents in a Protein Simple western blot imaging system.

## Supporting information

Supplementary Methods Table: Lysis buffer recipies

## Acknowledgments

The authors wish to thank Allison Maker and the Gumbiner lab for their assistance and use of their HPLC. The authors have no conflicts of interest to disclose. This work was supported by MH113540 (SEPS).

