## Supplementary figures and images for "Activity-dependent changes in synaptic protein complex composition are consistent in different detergents despite differential solubility"

### Figure S1

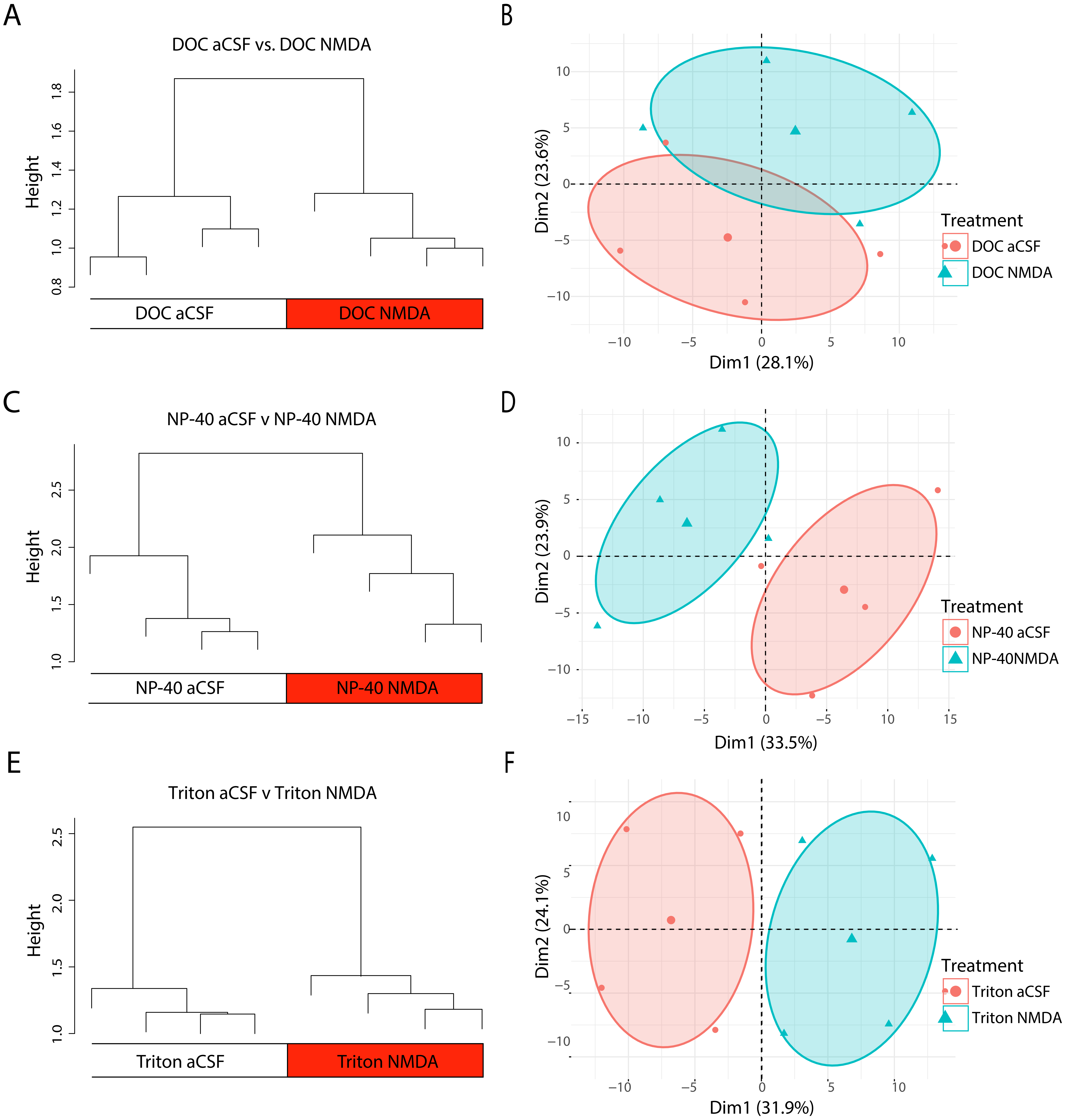
